# Topology of connective tissues: a key parameter in cellular heterogeneity, beyond composition and stiffness

**DOI:** 10.1101/2022.01.26.477902

**Authors:** Changchong Chen, Zeinab Ibrahim, Marion F. Marchand, Tristan Piolot, Sahil Kamboj, Franck Carreiras, Ayako Yamada, Marie-Claire Schanne-Klein, Yong Chen, Ambroise Lambert, Carole Aimé

## Abstract

Cellular plasticity is essential in physiological contexts, including pathological ones. It is the basis of morphogenesis and organogenesis, as well as tumorigenesis and metastasis. The extracellular matrix (ECM) is a key player in the generation of cellular heterogeneity. Advances in our understanding of cell plasticity rely on our ability to provide relevant *in vitro* models. This requires to catch the characteristics of the tissues that are essential for controlling cell fate. To do this, we must consider the diversity of tissues, the diversity of physiological contexts, and the constant remodeling of ECM along these processes. To this aim, we have fabricated a library of ECM models for reproducing the scaffold of connective tissues and basement membrane with different biofabrication routes based on the electrospining and drop casting of biopolymers. Using a combination of multiphoton imaging and nanoindentation, we show that we can vary independently protein composition, topology of connective tissues and stiffness of ECM models. Reproducing the features of a tissue and physiological context in turns allows to generate the complexity of the phenotypic landscape associated with the epithelial-to-mesenchymal transition (EMT) in human ovarian cancer. We show that EMT shift cannot be directly correlated with a unique ECM feature, which reflects the multidimensionality of living environments. Very importantly, our combinatorial approach allows us to provide *in vitro* models, where the impact of the topological cues on cellular phenotypes can be revealed, beyond protein composition and stiffness of the ECM matrix. On this line, this work is a further step towards the development of ECM models recapitulating the constantly remodeled scaffolding environment that cells face and provides new insights for the development of cell-free matrices.

## 1. Introduction

Cellular heterogeneity is the core of various cellular processes such as morphogenesis and tumor metastasis. During epithelial-to-mesenchymal transition (EMT), epithelial cells, organized in tissue with tight junctions and polarized morphology, can evolve towards less cohesive and highly motile cells. The associated high cellular heterogeneity, recently referred to as epithelial-to-mesenchymal plasticity (EMP), contributes to tumor heterogeneity.^1–3^ Moreover, cell heterogeneity has been correlated with chemoresistance.^4^ This is why regulating cellular heterogeneity is a significant challenge from a fundamental and therapeutic point of view. The extracellular matrix (ECM) plays a central role in the generation of cellular heterogeneity, being the scene of extensive transformations in diverse physiopathological or developmental contextes.^3–13^ Recent works have shed light on the importance of cell interactions with ECM components in cancer^14,15^ and during embryogenesis^16^ underscoring the regulation from both mechanical and biochemical stimuli. Yet, one of the largest hurdles to the better understanding of cellular heterogeneity is the establishment of models that recapitulate the heterogeneous and constantly evolving environment that cells explore. To address this combinatorial challenge, we use porous supports, called *patches* (Fig.1), to host ECM models while independently varying protein composition, ECM topology, and stiffness. (1) We exploit diversities in ECM proteins to mimic the scaffold of connective tissues (using type I collagen) and basement membrane (BM) (laminin and type IV collagen), which are both remodelled during cancer progression.^15^ (2) We uncouple protein composition, topology and stiffness by playing independently with the dimension of the patch and the processing of type I collagen (electrospinning (**es**) *vs* dropcasting (**dc**), Fig.1). (3) Very importantly, for all investigated ECM models, we reach physiological (including pathological) stiffness, which is a crucial prerequisite for such studies. (4) As cellular model, we use SKOV-3 human ovarian adenocarcinoma cells. This cell line displays large cellular heterogeneity during EMT.^17–20^ Indeed, ovarian cancers are very sensitive to their surrounding microenvironment. This includes the soluble environment - the ascite - which is an excess fluid in physio-pathological conditions^21^ playing a role in EMT,^20^ but also the biochemical and mechanical features of the ECM.

**Figure 1.**
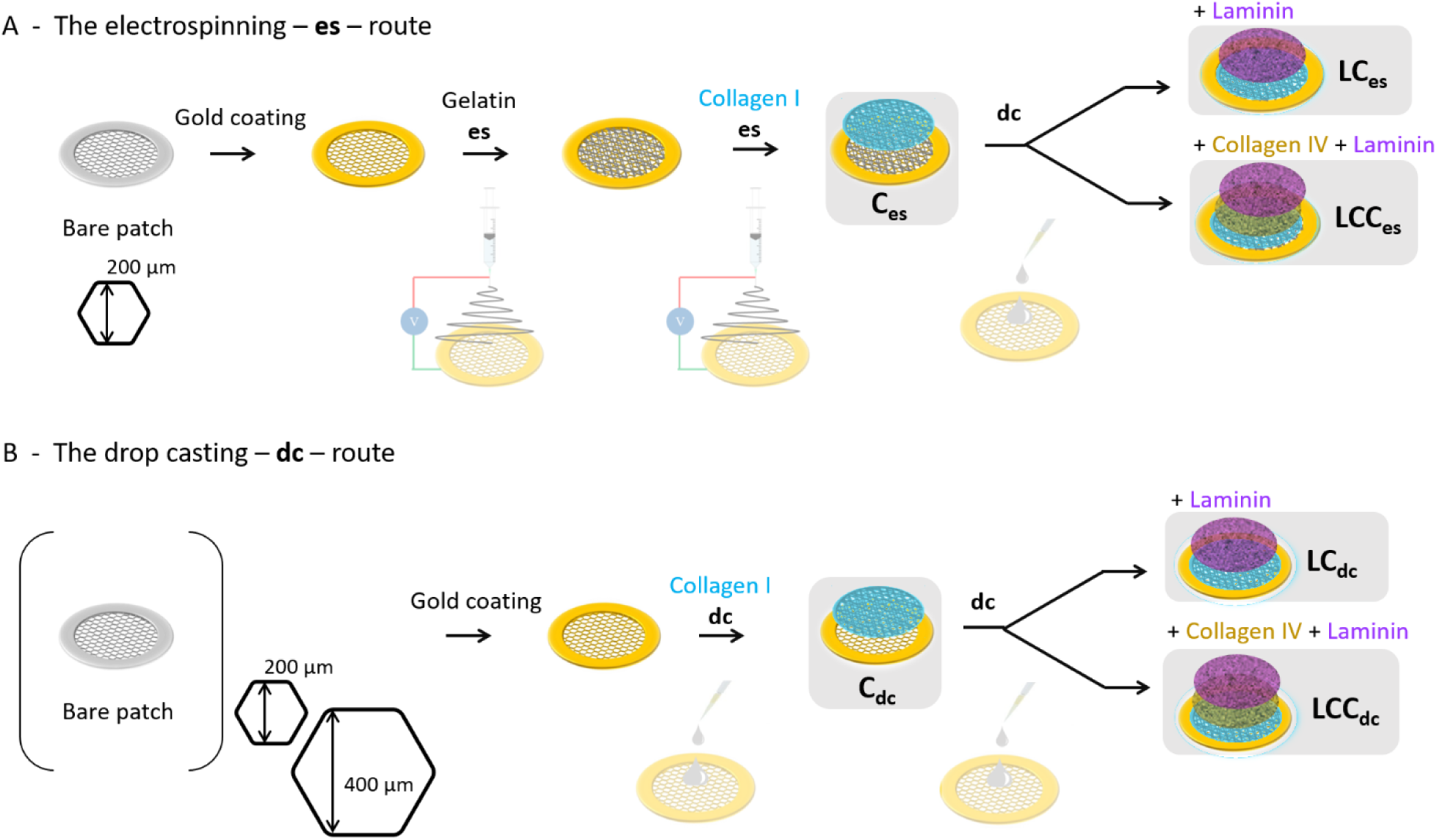
Scheme of the preparation of the ECM models based on (A) electrospinning (**es**) and (B) drop casting (**dc**) of ECM proteins including collagen I, collagen IV and laminin.

In a first step, we have used a combination of multiphoton microscopy and atomic force microscopy (AFM) to characterize the topology and mechanical properties of the ECM models. We have then explored the generated cellular heterogeneity in SKOV-3 cells depending on the features of the ECM models. Measurement of cell heterogeneity is crucial but remains difficult to quantify because of the existence of a continuum of phenotypic variations that cannot be carefully distinguished by biochemical bulk assays. To tackle this limit, we have combined qualitative immuno-fluorescence observations with a morphological single cell profiling quantification.^22^ We have focused on the variation of the nucleus structure, which is among the most important morphological changes observed in tumor cells.^23–25^ Indeed, quantitative analysis of cancer cell nuclear morphology has been used in cancer diagnosis. Here we have observed that the cellular heterogeneity generated by the diverse ECM models is represented by two shape modes: the size and shape, *i*.*e*. deformability. These cell morphological variations have a complex, not straightforward dependency on the features of the ECM models and cannot be solely attributed to variations in stiffness of the fabricated ECMs. This confirms that either of the parameters (protein composition, topology, stiffness) can be dominant for determining cell population features and for inducing EMT.^26^ Our combinatorial ability to provide ECM models is a very straightfoward tool for defining microenvironment conditions where one parameter can prevail over the other, while preserving biological relevance in the mean time. Very importantly, it provides a way to decorrelate connective tissue topology from protein composition and matrix stiffness. As such, this work will have a great impact for further biological studies and for improving the biofabrication of acellular matrices for clinical investigations.

## 2. Materials and methods

### 2.1 Type I collagen extraction and purification

Type I collagen was extracted and purified from rat tail tendons as previously described, except that we used 3 mM hydrochloric acid instead of 500 mM acetic acid.^27,28^ Collagen purity was assessed by electrophoresis and its concentration estimated by hydroxyproline titration.^29^ All other chemicals were purchased and used as received. Water was purified with a Direct-Q system (Millipore Co.).

### 2.2 Patch fabrication

The patches were fabricated by photolithography and soft-lithography. Firstly, a double layer SU-8 mold was fabricated by photolithography. The mesh layer was patterned on a silicon wafer using a 50 μm thick SU-8 negative photoresist by UV exposure at 250 mJ/cm^2^. Then, the honeycomb frame of 50 µm height was directly patterned on the mesh layer by another round of UV exposure at 250 mJ/cm^2^. After development in propylene glycol methyl ether acetate, this double layer SU-8 mold was then exposed in trimethylchlorosilane (TMCS, Sigma, France) vapor for 10 min. Afterwards, a mixture of PDMS (GE RTV 615) pre-polymer and its crosslinker at ratio of 10:1 (w/w) was casted on the SU-8 mold. After curing at 75°C for 4 hours, the PDMS layer was peeled off. Then, this replicated PDMS structure was placed on a glass plate and a solution of a photo-crosslinking polymer (Ormostamp®, micro resist technology) was injected in the free space of the PDMS-glass assembly, followed by UV exposure at 1500 mJ/cm^2^, and removal of the PDMS mold.

The patches were then coated with gold by sputter deposition for electrospinning (route A, Fig.1) or for improving hydrophilicity and drop casting (route B, Fig.1). For this, we used an Emitech K675X Sputter Coater System working at 125 mA for 30 seconds.

### 2.3 *Electrospinning (*es)

The gelatin solution was prepared by dissolving 100 mg of gelatin in a mixture of 420 μL acetic acid, 280 μL ethyl acetate and 200 μL DI water. Citric acid (10 mg) was then added as thermal cross-linking agent before **es**. The mixture was stirred for 4 hours at room temperature. Gelatin was electrospun onto a gold-coated patch fixed on a tin foil (7 cm in diameter) at a voltage of 11 kV for 7 minutes at a flow rate of 0.2 mL/h, with a controlled humidity of 35% at room temperature (20~25°C). The distance between the metal needle and the patch was 9-12 cm. After **es**, the patch was detached from the tin foil and transferred into an oven working at 140°C for 4 hours for high temperature cross-linking of gelatin.

Type I collagen solution (1.7 mg.mL^−1^ in 30 mM HCl and 75 vol% ethanol) was electrospun for 1 hour onto gelatin fibers support at 11 kV with a flow rate of 0.2 mL/h, with a controlled humidity of 25% at room temperature (20~25°C). The distance between the metal needle and the counter electrode was 11 cm. After **es**, collagen fibrillogenesis was induced in vapor phase by placing the collagen-coated patches in a chamber saturated with ammonia overnight.

### 2.4 *Drop casting (*dc)

Type I collagen (0.5 mg/mL in PBS, pH=9.0), laminin (10 μg/mL) and type IV collagen (0.5 mg/mL) were drop cast by pouring 20 µL of the solution on the patch and dried in air at room temperature.

### 2.5 SEM imaging

Samples were coated with gold for 60 s with a sputtering current of 50 mA before imaging. Samples were fixed on conductive-tapes for imaging with a TM3030 Tabletop Microscope (Hitachi High-Technologies Corporation, Japan) equipped with TM3030 software and working at an acceleration voltage of 15 kV.

### 2.6 Second harmonic generation / 2-photon excited fluorescence

We used a custom-built laser-scanning multiphoton microscope and recorded second harmonic generation (SHG) and 2-photon excited fluorescence (2PEF) images in parallel as previously described.^30^ Excitation was provided by a femtosecond titanium–sapphire laser (Mai-Tai, Spectra-Physics) tuned to 860 nm, scanned in the XY directions using galvanometric mirrors and focused using a 25× objective lens (XLPLN25X-WMP2, Olympus), with a resolution of 0.35 μm (lateral) × 1.2 μm (axial) and a Z-step of 0.5 µm for the acquisition of Z-stack images. We used circular polarization in order to image all structures independently of their orientation in the image plane, using 100 kHz acquisition rate and 420×420 nm^2^ pixel size.

Patches were observed in duplicate to check for reproducibility and for each patch, three different areas were imaged to verify the homogeneity of the biopolymer coating.

### 2.7 AFM nanoindentation

AFM experiments were performed on hydrated ECM-models in cell culture medium at room temperature using a NanoWizard 4 (JPK BioAFM, Berlin, Germany) mounted on an Axio Observer microscope (Zeiss, Oberkochen, Germany) placed on a vibration isolation table. Before each experiment, the cantilever spring constant was accurately determined upon calibration in cell culture medium by the thermal noise method.

Young’s moduli were determined by colloidal probe force spectroscopy using a gold coated cantilever (0.01 N/m) equipped with a 6.44 μm bead probe (NanoAndMore, Paris, France). Approach and retraction speeds were kept constant at 5 µm/s, ramping the cantilever by 10 µm with a 0.4 nN threshold in a closed z loop. In this option, the feedback system readjusts the initial piezo-position for each force–displacement ramp so that the maximal force applied to the sample remains constant.

AFM force-distance curves were transformed to force-indentation curves and fitted using the JPK data processing software. Curves were fitted using the contact-point independent linear Hertz-Sneddon model. The Hertz model assumes infinite sample thickness, which was approximated by using small indentation (typical indentation depth 500 nm). Automated curve fitting was applied using fitting range of 100% of curve. Force-distance curves were measured in at least 8 positions for 3 different areas of each ECM model.

Statistical analysis was performed using Origin software. Analysis of variance (ANOVA) with the Tukey’s Multiple Comparison test was used for all multiple group experiments. P values < 0.05 were deemed significant. Values in graphs are the mean and standard error of mean (*p < 0.05, **p < 0.001, ***p < 0.0001).

### 2.8 Cell culture experiments

SKOV-3 cells (ATCC1, HTB77™), human ovarian adenocarcinoma cell line, were purchased from ATCC (American Type Culture Collection, Manassas, VA). SKOV-3 cells were cultured in RPMI-1640 glutaMAX containing 0.07 % (v/v) sodium bicarbonate supplemented with 10% fetal calf serum and 1% (v/v) penicillin streptomycin (all reagents were purchased from Thermo Fisher Scientific). Cells were cultured in T25 cell culture flasks in a humidified air atmosphere with 5% CO_2_ at 37°C.

ECM models were immersed in 70% ethanol for 5 min for sterilization, and washed 3 times for 5 min in sterile PBS. Cells were seeded at a density of 30 000 per patch. Patches immersed in cell culture medium were then transferred to a controlled atmosphere (37°C, 5% CO_2_) for 1 day.

Experiments were run in triplicate for each ECM condition.

### 2.9 Immunofluorescence

Cells were fixed in 4% paraformaldehyde (PFA) in PBS for 10 minutes, rinsed three times with PBS. The cells were permeabilized with 0.5% Triton X-100 in PBS, washed again and saturated with PBS containing 3% BSA for 30 min. Cells were incubated overnight at 4°C with Alexa Fluor 543-conjugated vimentin antibody (ab202504, Abcam®) at a 1/1000 dilution. After washing, the actin cytoskeleton was stained with Alexa Fluor 488 Phalloidin in PBS (containing 1% DMSO from the original stock solution, Abcam®) for 40 min at room temperature in a dark chamber. Cell nuclei were then stained with DAPI (4,6-diamidino-2-phenylindole dihydrochloride, Molecular Probe®) for 15 min. Immunofluorescent labelling was observed with a confocal microscope (LSM710, Zeiss) equipped with 405 (DAPI), 488 (phalloidin), and 543 (anti-vimentin antibody) nm lasers and with LSM ZEN 2009 software. We used 1 μm z-stack intervals and sequential scanning for each wavelength. Images were processed with ImageJ.

For the characterization of the basement membrane by 2PEF, ECM models were stained with Alexa Fluor 488^®^-conjugated collagen IV monoclonal antibody (eBioscience™, 53-9871-82) incubated overnight at 4°C or 2 h at room temperature at a concentration of 10 μg.mL^−1^, together with laminin polyclonal primary antibody (Thermo Fisher Scientific, PA1-16730) overnight at 4°C or 2 h at room temperature at a concentration of 1 μg.ml^−1^, followed by 1 to 2 h incubation at room temperature with a goat–anti rabbit-Alexa Fluor 610 secondary antibody at a dilution of 1/250.

### 2.11 Statistical analysis of cell morphology

Cells were analyzed by principal component analysis of nucleus morphology.^31,32^ We used Celltool^33^ to identify single cells based on their nuclei. Experiments were run in triplicate for each ECM condition. The analysis of the morphology of the nucleus was carried out on more than 300 cells per condition, *i*.*e*. on the triplicate, by considering each cell data as independent. This allows good sampling of the statistical distribution associated with each microenvironment.

## 3. Biofabrication of ECM models, a combinatorial challenge

The ECM can be described in terms of protein composition, organization and of the resulting mechanical properties such as stiffness. A first concern to reach physiological stiffness, is that self-supported ECM models should be favored to limit the contribution of the support. This can become challenging when attempting to get biologically relevant thin tissue models. To this aim, we have microfabricated *patch* supports with a honeycomb pattern (Fig.2A,B), which allows to combine the standing of the ECM model while leaving large areas of tissues unsupported. Besides, dimensions can be tuned using photo-lithography processes to accommodate in the meantime constraints associated to tissue processing, adjustable mechanical properties, cell culturing and associated monitoring. One other key advantage of this support compared to conventional solid substrate is its high porosity allowing to preserve the porous features of connective tissues and BM composing the ECM. These patches have been shown to successfully allow the maintenance^34^ and differentiation of human induced pluripotent stem cells (hiPSCs) into motor neurons^35^ and functional cardiomyocytes,^36^ and have been used for various bioassays further underscoring their easy integration into a diversity of devices.^37–40^ In this work, for the elaboration of ECM models, circular patches are used having a diameter of 1.1 cm (Fig.2A) with variable dimensions of the honeycomb pattern (Fig.2B yellow arrow). Two dimensions of honeycombs (200 and 400 µm) were tested for the **dc** route to improve the self-standing features of the ECM models and limit the contribution of the support. Note that for the **es** route, only 200 µm patches were used due to the small size of electrospun collagen fibers (see below).

**Figure 2.**
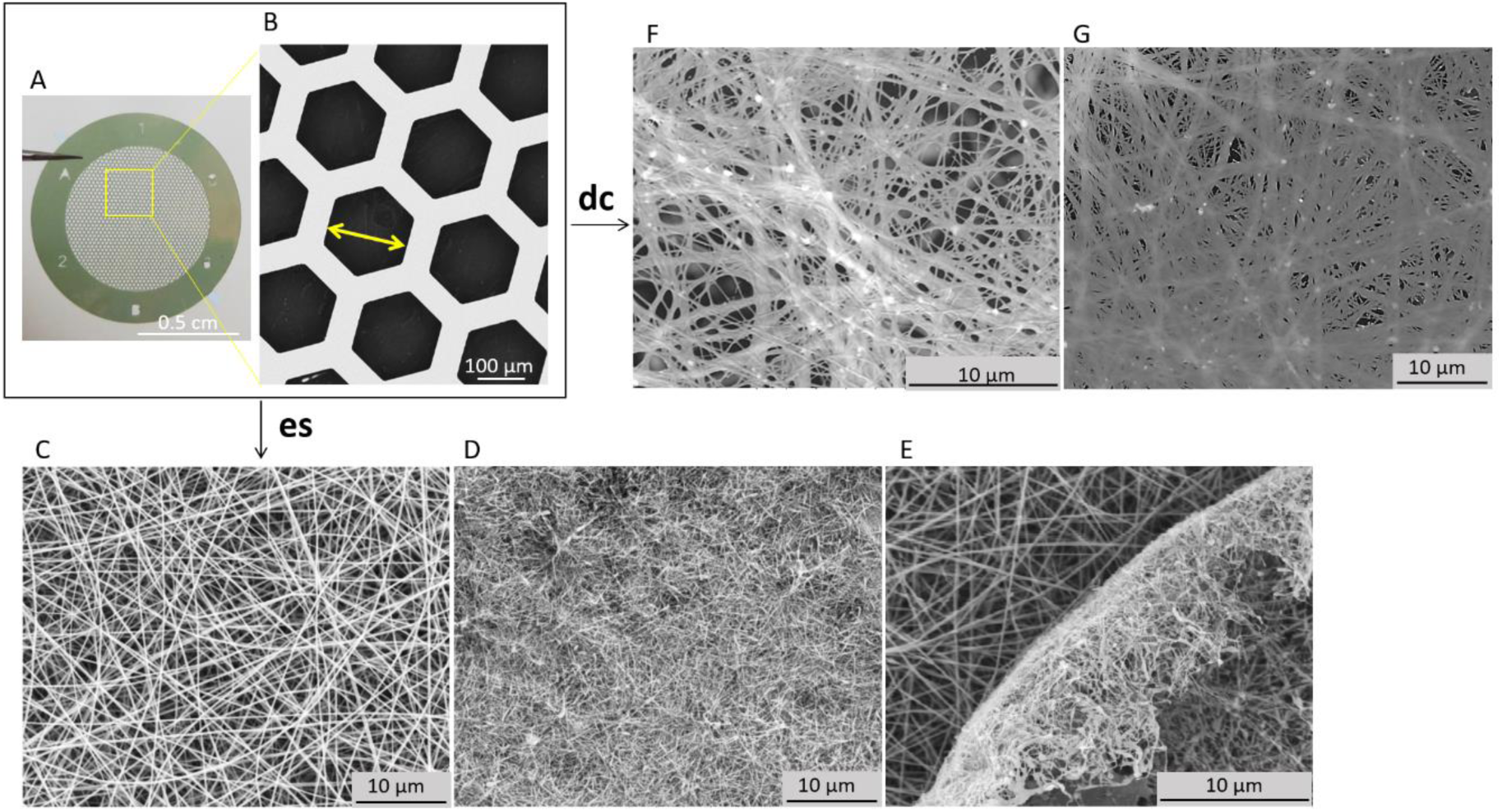
(A) Photograph and (B) SEM image of the microfabricated bare patch. SEM images of patch after electrospinning (**es**) of (C) gelatin, and (D,E) electrospun collagen I on top of the gelatin layer (the image in (E) shows a default on the border of the patch clearly showing the layered structure), and after dropcasting (**dc**) collagen I on (F,G) 200 and 400-µm patch respectively.

A second key point for mimicking living microenvironments is that composition, organization and stiffness are all constantly evolving during physiological and pathological processes.^5,41^ We have selected three ECM proteins for their contribution to cell invasion,^42–44^ and for reconstructing both connective tissues – type I collagen – and BM – laminin and type IV collagen. Independently from the composition, the organization of the connective tissue models was varied using two different processes for type I collagen deposition: electrospinning (**es**) and drop casting (**dc**). Electrospinning provides a net-like structure made of cross-linked nanofibers and is thus relevant to mimic the complex *in vivo* ECM organization. The process consists in the extrusion of a soluble biopolymer assisted with electrical field. This first requires to coat patch with conductive gold to accumulate collagen fibers on the support (Fig.1A). It has been shown that **es** preserves the native state of collagen I, producing small fibers (*ca*. 60 nm in diameter).^45^ Because the collagen nanofibers are small with respect to the dimension of the honeycomb pattern (200 µm-wide honeycombs, see the yellow arrow Fig.2B), we first deposited a layer of electrospun gelatin (Fig.2C) and did not further increase the size of the honeycomb frame. Given the viscoelastic properties of gelatin, micrometric fibers can be obtained by **es**. This layer in turn successfully supports the electrospun collagen on the patch (Fig.2D), while preserving the porosity of the support. This double processing ends with a layered structure of nanofibers (Fig.2E).

Two routes are thus compared to obtain ECM models: (A) the **es** route and (B) the **dc** route (Fig.1). The **es** route leads to the formation of electrospun collagen I nanofibers (**C**_**es**_), eventually topped with a cast layer of laminin (**LC**_**es**_) or with successive layers of collagen IV and laminin (**LCC**_**es**_). Alternatively, in the drop casting route, collagen I is drop cast onto the patch (**C**_**dc**_), and eventually topped with a layer of laminin (**LC**_**dc**_) or with successive layers of collagen IV and laminin (**LCC**_**dc**_). Note that before drop casting, patches were coated with gold to keep the mechanical properties of the frame constant with respect to the **es** conditions, and to enhance hydrophilicity of the frame for drop casting.

## 4. Features of ECM models

### 4.1 Topology

The nine different ECM models were characterized by multiphoton microscopy combining second harmonic generation (SHG) and 2-photon excited fluorescence (2PEF). This multimodal imaging allows to monitor in parallel unlabeled collagen I with high specificity towards its fibrillary hierarchical organization through SHG, and collagen IV and laminin after immunostaining using two 2PEF channels. The reproducibility and homogeneity of the biopolymer coating were checked by imaging three different areas for each condition, and prepared as patch duplicate (in total 6 areas imaged per condition). Figure 3 presents a gallery of SHG/2PEF images with top, bottom and side views for each ECM model.

**Figure 3.**
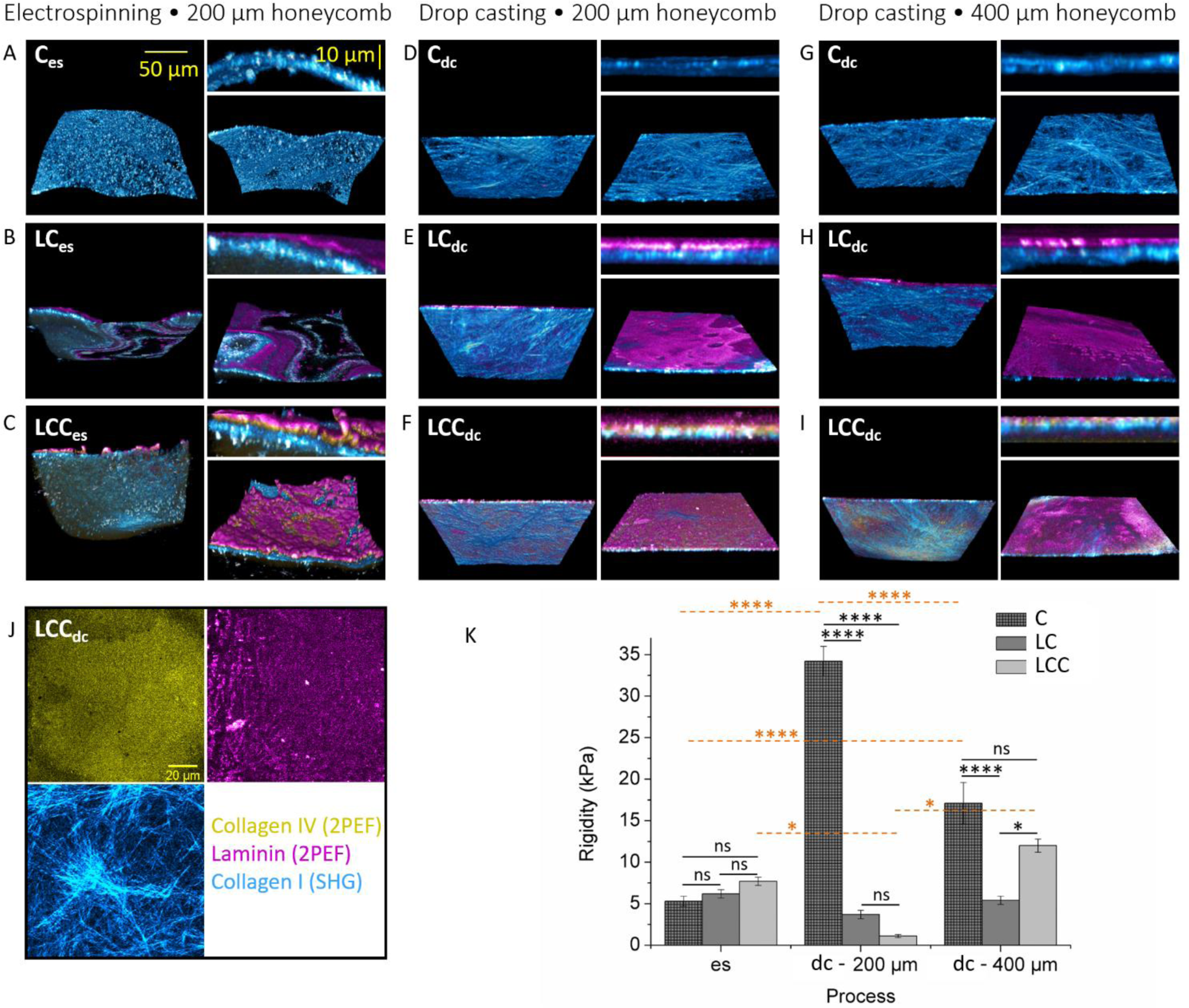
SHG (collagen I in cyan) and 2PEF (laminin in magenta and collagen IV in yellow) images of the ECM models after (A-C) **es** and **dr** on (D-F) 200 µm- and (G-I) 400 µm-honeycombs. For each sample, the left image is the bottom view, the top right one the side view, and the bottom right image is the top view. (J) Details of the single channels recorded for multiphoton characterization of the **LCC**_**dc**_ sample: top left is the 2PEF channel for anti-collagen IV antibody detection (in yellow), top right is the 2PEF channel for the anti-laminin antibody detection (in magenta) and the bottom left is the SHG channel for unstained collagen I detection (in cyan). (K) Mean Young’s modulus values for all ECM models measured by peak force tapping AFM-nanoindentation. In the plots, the column represents Mean with SEM. (*p < 0.05, ***p < 0.00001, ns for non-significant; using Tukey’s Multiple Comparison test).

All scaffolds share common features including a thickness of a couple of microns, and a layer-by-layer structure: for all LC and LCC scaffolds, the layer structure is confirmed with a bottom collagen I layer (cyan in Fig.3) mimicking connective tissues and a top layer of laminin (magenta in Fig.3) with/without collagen IV (yellow in Fig.3). Side views in Figure 3C,F,I better reveal the layer-by-layer structure in LCC scaffolds as collagen IV signal is difficult to identify on 3D reconstructions due to its embedding in between collagen I and laminin. The presence of collagen IV was further evidenced by the single 2PEF channel image as shown for **LCC**_**dc**_ (Fig.3J and see also Supporting Information for the full characterization of all the ECM models).

The scaffolds obtained by the **dc** route, whatever the dimension of honeycombs, are qualitatively similar with flat, layered structures based on a bottom fibrillar network of collagen I (Fig.3D-I). Topologic variations specific to the process can be observed with electrospun scaffolds: (1) **es** scaffolds are not as planar as the drop cast ones. This is attributed to the heterogeneous distribution of the electric field used for **es** throughout the patch, *i*.*e*. on the gold frame *vs* in empty honeycombs. Note that in those images, the underlying gelatin fibrillary layer observed by SEM (Fig.2D,E) is not stained. (2) The structure of the collagen I network is very specific to the process, *i*.*e*. **es** *vs* **dc**, as shown in particular with pure collagen I scaffolds (see **C**_**es**_ *vs* **C**_**dc**_ scaffolds). The **es** scaffolds exhibited a grainy-like pattern with alternating high and low SHG intensity. As it has been reported earlier, this signs for the small size of the fibrils with high entanglement.^45^ Indeed, given that the diameter of the fibrils (*ca*. 60 nm) is small relatively to the focal volume (0.35 μm (lateral) × 1.2 μm (axial)), many fibrils may be simultaneously observed in one pixel. At the intersection of these fibrils, a centrosymmetric distribution of the molecules is obtained in the focal volume, which leads to a local decrease of SHG signal.^45^ If the network is highly entangled, SHG decrease is widespread throughout the focal volume leading to a grainy-like image (Fig.3A). In contrast, after drop casting, collagen I networks are made up of large entangled fibrils as observed in Figure 3D and G.^46^

### 4.2 Mechanical properties

Nanoindentation was implemented by peak force tapping AFM in liquid environment, which is particularly powerful for characterizing soft biomaterials with nanometer depth and picoNewton force resolution.^47–52^ To measure quantitatively the mechanical heterogeneity of ECM stiffness, the median of Young’s modulus values was measured in at least 8 positions for 3 different areas for each type of scaffold. Here, we observed median Young’s modulus values in the 1 to 35 kPa range (Fig. 3K and Table S1). The lower so-called “soft” values in the kPa order are similar to what is measured from physiological ECM, while the higher “stiff” values with tens of kPa have been associated with severe diseases such as cancer and fibrosis.^53–55^ **es** scaffolds were found to be soft and homogeneous (see Table S1). Non significant variations in Young’s moduli were measured after addition of laminin and collagen IV (5.3 to 6.2 and 7.7 kPa). In contrast, after drop casting of collagen I, stiff scaffolds were obtained (34.2 and 17.1 kPa on 200 and 400-µm patch respectively) and exhibited higher heterogeneity than **es** scaffolds. This is attributed to the formation of the large bundles of drop cast collagen I as observed by SHG that accounts for both higher stiffness and higher local heterogeneity when compared to the more homogeneous networks of small collagen fibers obtained by **es**. The contribution of the underlying gelatin layer on the stiffness of **es** scaffolds was ruled out by drop casting collagen I on a gelatin layer, resulting in Young’s modulus of 35.8 kPa (Table S1). This value is similar to the drop cast collagen I in absence of gelatin fiber layer. This indicates that beyond the presence of the gelatin layer, and for indentation depth of the scale of indentation by the cell (500 nm), the choice of **es** *vs* **dc** dominates the resulting topology and stiffness of ECM models.

For drop cast scaffolds, the addition of laminin or collagen IV-laminin, softens significantly the scaffold, with Young’s modulus values droping from 34.2 to 3.7 and 1.1 kPa respectively for 200 µm-wide honeycombs and from 17.1 to 5.4 and 12 kPa for 400 µm-wide honeycombs. In addition to their softening, the ECM scaffolds appear much more homogeneous in terms of stiffness distribution. This is attributed to the formation of a homogeneous layer of small structures as identified by 2PEF, compared to the large collagen bundles associated to a high stiffness and heterogeneity obtained by **dc**.

Let us now take a closer look at the stiffness measured for a given protein composition, when varying the process and patch dimensions (in orange in Fig.3K; for clarity, only significant differences have been marked). No significant difference in Young’s moduli were measured between **LC** scaffolds as compared to **LCC**. In addition, the most important differences in Young’s moduli were observed for pure collagen I scaffolds, the **es** one being the softest. Changing process leads to different protein organizations and resulting ECM topology, as confirmed by SHG (Fig.3A,D,G). The access to these different topologies is biologically very relevant as it reflects the diversity in organization and fiber size of the different living tissues. Moreover, significant differences were measured for drop cast collagen on 200 and 400 µm pattern going respectively from 34.2 to 17.1 kPa. This shows that playing with the dimensions of microfabricated porous architectures is another way for imparting a mechanical response to self-standing ECM models.

## 5. Impact of the ECM model on ovarian cancer cells

The possibility to generate a phenotypic heterogeneity and reproduce the EMP spectrum has been assessed on the different types of patches. Human ovarian adenocarcinoma SKOV-3 cell line has been selected for modelling cellular heterogeneity because their phenotypes *in vitro* are representative of the heterogeneity observed in ovarian cancer.^17–20^

The evolution of cell phenotype on the different ECM models was monitored after one day by immunofluorescence. Experiments were run in triplicate and nucleus morphology analysis was carried out on more than 300 cells per condition. Cells were stained for the nucleus (DAPI) and for the cytoskeleton by combining staining of actin (phalloidin) and of vimentin (anti-vimentin antibody). Vimentin is a cytoskeletal protein upregulated during cell transitioning events commonly used as a marker of EMT, and whose expression is associated with mesenchymal phenotypes. Confocal microscopy observations revealed important phenotypic variations depending on the ECM models. Two extreme phenotypes can be described, together with a multiplicity of intermediary states along the EMT phenotypic continuum. First, cells with a compact and round nucleus were identified, which is characteristic of most epithelial cells (Fig. 4A). This is also associated with a perinuclear distribution of vimentin as clearly identified in Figure 4A4. On the other side of the phenotypic continuum, elongated cells having a spindle-like morphology were also observed (Fig. 4B). This cells exhibited an elongated actin cytoskeleton. Remarkably the cortical distribution of vimentin is observed, which drastically differs from the epithelial-type of cells described previously. In that case, the vimentin cytoskeleton co-localizes with the actin one (Fig.4B4). This is characteristic of mesenchymal-type cells. This illustrates the high level of reorganization of cells and shows the influence of the ECM microenvironment on cytoskeleton reorganization. Importantly, it also confirms the possibility to generate cellular heterogeneity in ovarian cancer cells based on the alteration of the microenvironment, as previously reported from circulating ascites.^20^

**Figure 4.**
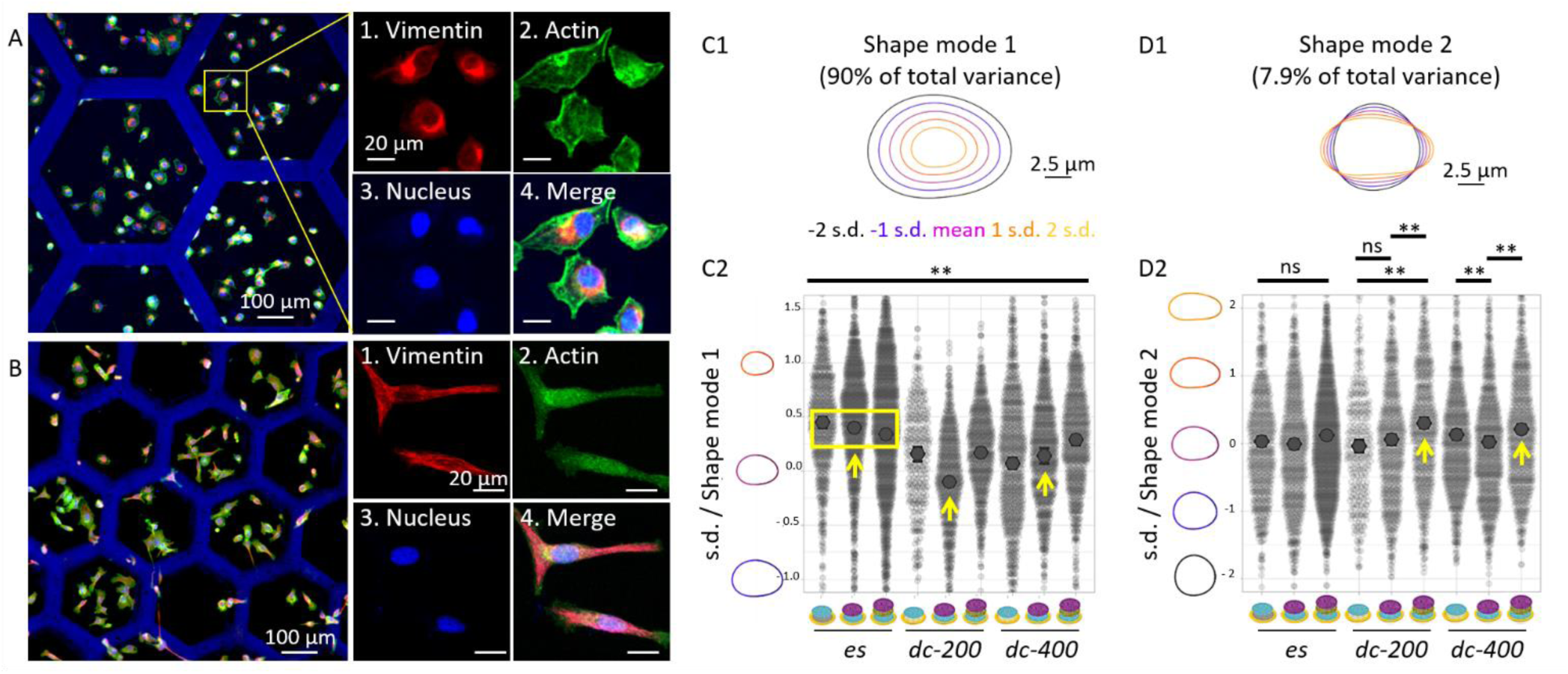
Immunofluorescence images of SKOV-3 cells one day after seeding on (A) **400-C**_**dc**_ and (B) **200-LCC**_**dc**_. Cells were stained for (1) vimentin (red), (2) actin (green), and (3) nuclei (blue); (4) is the merge of the three. The scale bar is 20 µm for all zoom-in images. (C-D) Principal component analysis of nucleus morphology one day after seeding on the different ECM models: morphological variance for (C1-2) shape mode 1 – nucleus size mostly – and (D1-2) shape mode 2 – nucleus shape and deviation from sphericity mostly (comparison is between all samples for C2 and between samples at extremities of bar in D2, with n.s: non significant; *p<0.05; **p<0.001). The color code is common to C1 and D1. All plots were realized using SuperPlotsOfData.^56^

We then quantified these changes in morphology at the cell level by automated object segmentation and measurement using principal component analysis (PCA), over 10000 cells.^31,32^ We focused on shape metrics because it is less prone to external contributions when compared to the analysis of fluorescence intensity of specific markers (staining, acquisition conditions). We identified cells based on their nuclei, since nuclei are mechanosensors of external ECM cues.^22^ Based on the segmentation of nuclei, two parameters, called *modes*, were found to recapitulate the cellular heterogeneity generated on the ECM models. Figures 4C1 and D1 show the two corresponding histograms that represent the diversity spectrum of all segmented nuclei (the color code is common to C1 and D1). Shape mode 1 mostly accounts for the size of the nucleus and represents 90% of total variance (Fig.4C1). Shape mode 2 represents nucleus shape, *i*.*e*. mainly deformation correlated with EMT shift. It accounts for 7.9% of variability (Fig.4D1). The corresponding mean values and distributions of nucleus size and shape as a function of ECM models are further detailed in Figures 4C2 and D2 respectively.

Let us first examine the heterogeneity in nucleus size. Figure 4C2 shows that nucleus sizes measured over more than 300 cells per condition were found to be significantly different when comparing all ECM models with each other. Remarkably, no specific trend could be identified with respect to the protein composition within each of the three groups of scaffolds. A small increase in size was observed from **C**_**es**_ to **LC**_**es**_ and to **LCC**_**es**_. The inverse trend was observed for 400-drop cast samples and no common features identified with 200-drop cast samples. In contrast, significant trends were observed when comparing **es** *vs* **dc** routes. Cells on **es** scaffolds (yellow box, Fig.4C2) displayed compact nuclei whatever the protein composition, when compared to cells seeded on drop cast scaffolds with either 200 or 400 µm dimensions. In these later cases, much larger nuclei were measured. This is an interesting result given that the most important difference when comparing the three groups is the topology of reconstructed connective tissues from collagen I (see SHG in Fig.3A-I) rather than the stiffness (except for the pure **C**_**es/dc**_). The importance of topology over stiffness is also illustrated when comparing **LC**_**es**_, **200-LC**_**dc**_ and **400-LC**_**dc**_ that exhibit similar stiffness, protein composition but different connective tissues topology resulting in important variations of the nucleus size (yellow arrows, Fig.4C2).

Changes in shape mode 2 mostly account for deformation of nucleus correlated with EMT. In that case, non significant nucleus deformations were measured for **es** scaffolds, and when comparing **200-C**_**dc**_ and **200-LC**_**dc**_ (Fig.4D2). The most important deformations of the nucleus were measured on **LCC**_**dc**_ models, *i*.*e*. for a given topology of collagen I but also for a given protein composition, independently of the stiffness of the ECM model (yellow arrows, Fig.4D2).

## 6. Discussion

Because cells respond to their microenvironment, including mechanical and biochemical cues from the ECM, it is important to discriminate protein composition, topology and ECM stiffness to deepen our understanding and improve the biofabrication of relevant *in vitro* models. In addition, the ECM is constantly remodeled during development and disease, as previously described by SHG in the context of ovarian cancer.^57–59^ This makes it crucial to have the ability to adjust ECM models to meet different requirements.^60^ Here we have elaborated a library of ECM models, where not only stiffness, but also ECM composition and the topology of connective tissues could be varied playing with the processing of biomolecules (**es** *vs* **dc**) and the dimensions of microfabricated supports. Using SHG and AFM, we have characterized the composition, topology and stiffness of nine different ECM models that all reproduce the native topology (SHG/2PEF) and stiffness (nanoindentation) of physiological – including pathological – ECMs. For the biofabrication of connective tissues scaffold, we have shown that the method used to process type I collagen dominates the topology and associated stiffness of the ECM model. Interestingly, from these connective tissue models, the addition of an artificial BM was found to soften the ECM, and this independently from collagen I process.

SKOV-3 cellular heterogeneity on those ECM models was quantified through two modes defining the evolution of the size and shape – deformation – of the nucleus. Two important results to emphasize here are (1) a clear difference in cell morphology measured on different ECM models, and (2) the absence of a direct trend between nucleus size/deformation and protein composition, topology or/and Young’s modulus. This highlights the multidimensional correlation of the different biophysical and biochemical features that cannot be restricted to one varying parameter, not even the stiffness.^26,43^ This can explain the controversy found in litterature on the dependency of SKOV-3 cell fate with stiffness. SKOV-3 cells are clinically defined as having an epithelial morphology.^61^ Different groups have reported that soft matrices promote EMT and the acquisition of mesenchymal spindle-like cell shape together with the increase in vimentin expression.^62,63^ In contradiction, others have shown that increasing stiffness is associated with cell spreading and migration.^64^ All these works concern the same cell line and stiffness range but with no concern on protein nature and topology. Our strength in providing combinatorial ECM models is that it allows to discuss protein composition, topology and stiffness in a single study in which the systems are directly comparable, which cannot be achieved when comparing results coming from different strategies and cellular models. More importantly, it provides ECM models that may be more prone to mimic a given tissue and different physiological contexts, where either mechanical cues, molecular recognition or topological cues may prevail.

Importantly, this work shows that the cells are very sensitive to the topology of the collagen I matrix, which is a key constituent of the connective tissue scaffold (see for example nucleus size measured on **es** *vs* **dc** scaffolds). This parameter has been quite underestimated for years. In particular here, SHG/2PEF has revealed that the **dc** models are much more planar than the **es** one, whose curvotaxis has been shown to affect cell-ECM interactions.^65^ In addition, collagen I processing allows controlling the size of collagen bundles and the resulting heterogeneity in stiffness over the ECM models. The ability to control collagen organization is highly relevant for improving *in vitro* models as collagen topology varies from tissue to tissue providing them with their function.^66,67^ As such, and as revealed here, this should systematically be taken into account when biofabricating tissue models.

Interestingly, both metrics – size and shape of nuclei – were found not to evolve concommitantly for all ECM models. This is remarkable *per se* and may be attributed to the existence of different mechanotransduction pathways^6^ differently balanced over the ECM models. Moreover, the different patterns of evolution between shape mode 1 and 2 are probably at the origin of the diversity of the landscape characteristic of EMT plasticity, where different subpopulations of tumor cells with hybrid phenotypes were identified.^68^ Ultimately, our strategy offers the possibility to generate a broad diversity of phenotypic variations to be isolated on defined ECM models that take into account the biochemistry, mechanical properties, and topology of the microenvironment for further mechanistic investigations.

## 7. Conclusion

We have built *in vitro* models that recapitulate the biochemical, mechanical and topological characteristics of the cell microenvironment. ECM models reproducing both the scaffold of connective tissues and basement membrane allow to investigate the interplay between protein composition, topology and stiffness in regulating cellular heterogeneity in SKOV-3 cells. These cellular changes in response to ECM cues resemble many morphological features associated with EMT. This generates new knowledge on cell heterogeneity during EMT, where the topology of connective tissues was found to impact nucleus size and shape independently of stiffness and protein composition. This is particularly important given that more than contributing to cancer progression, EMT plasticity is also implicated in chemotherapy resistance. As such, our results have an important dual impact in translational research: (1) they promote the design of new devices for organoid culture needed to model human development and disease, and (2) they will improve commercially available acellular matrices that are currently used in various clinical settings, introducing signals from the microenvironment.

## Acknowledgements

We thank the China Scholarship Council for the PhD grant of Changchong Chen, Christophe Hélary and Gervaise Mosser for their help in the extraction and purification of type I collagen, and Clothilde Raoux for her constant support with SHG/2PEF experiments. This project has received financial support from the CNRS through the MITI interdisciplinary programs. Multiphoton imaging at LOB was partly supported by the Agence Nationale de la Recherche (contract ANR-11-EQPX-0029 Morphoscope2). This work benefited from the technical contribution of the Institut Pierre-Gilles de Gennes joint service unit CNRS UAR 3750. The authors would like to thank the engineers of this unit for their advice during the development of the experiments: Bertrand Cinquin, Nhung Dinh, Audric Jan, Kévin Phan. We thank members of MEC-uP Team for helpful discussions and comments. Johanne Leroy-Dudal and Rémy Agniel for experimental discussions. This work was funded by CY Initiative of Excellence (grant “Investissements d’Avenir” ANR-16-IDEX-0008).

