## Supplementary materials for "Topology of connective tissues: a key parameter in cellular heterogeneity, beyond composition and stiffness"

### Supporting Information

#### 1. Features of ECM models

*1.1 Second harmonic generation / 2-photon excited fluorescence*

*1.2 AFM nanoindentation*

### 1. Features of ECM models

#### *1.1 Second harmonic generation / 2-photon excited fluorescence*

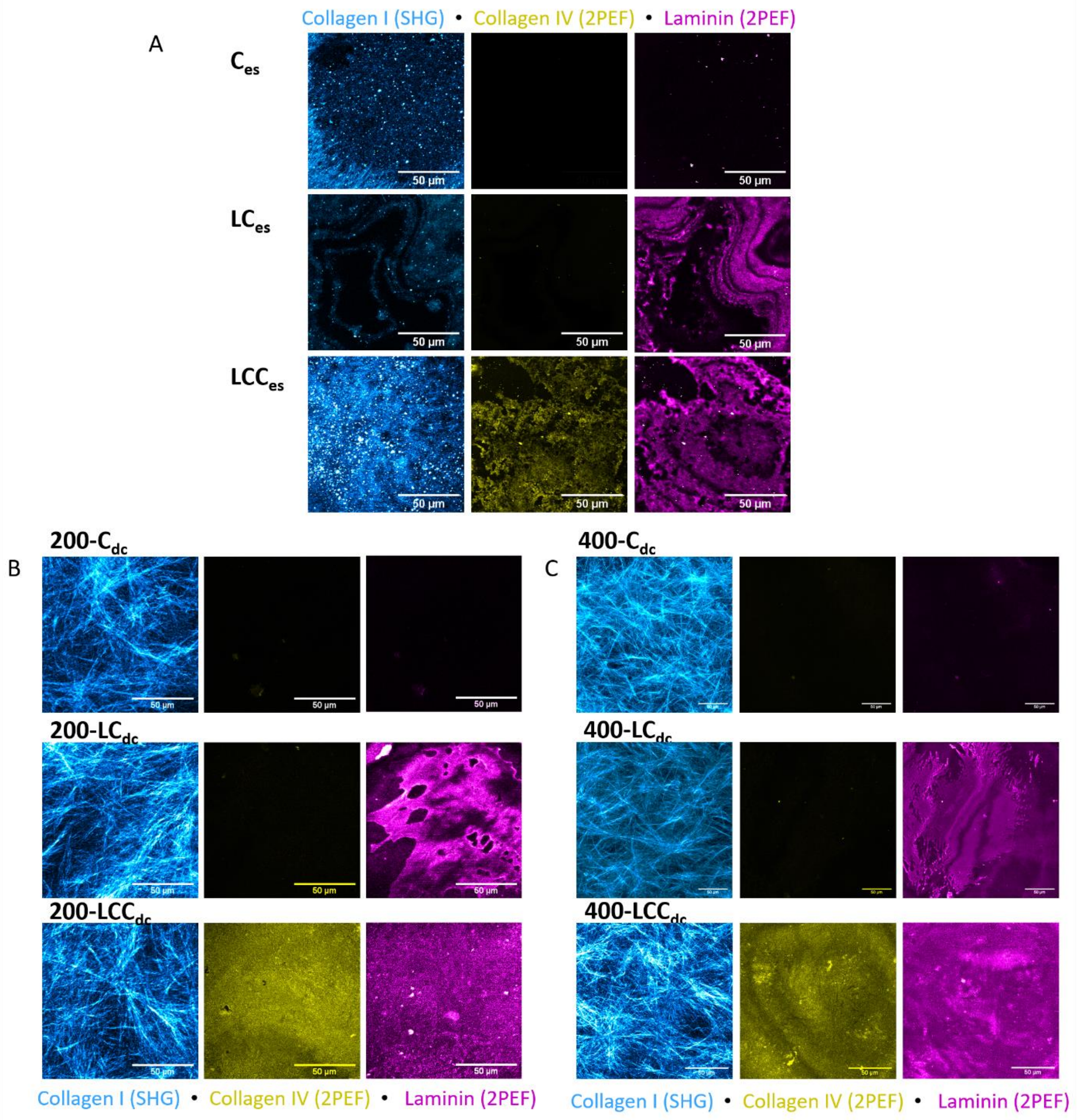


**Figure S1:** SHG (endogeneous signal of collagen I in cyan) and 2PEF (laminin in magenta and collagen IV in yellow) images of the ECM models after (A) **es**, and **dc** on (B) 200 µm- and (C) 400 µm-honeycombs. Patches were observed in duplicate to check for reproducibility and for each patch, three different areas were imaged to verify the homogeneity of the biopolymer coating.

*1.2 AFM nanoindentation*

| | **Mean** (kPa) | **Standard Deviation** | **SE of Mean** |
|---|---|---|---|
| **$C_{es}$** | 5.3 | 2.7 | 0.6 |
| **$LC_{es}$** | 6.2 | 2.5 | 0.5 |
| **$LCC_{es}$** | 7.7 | 2.6 | 0.5 |
| **200-$C_{dc}$** | 34.2 | 12.2 | 1.8 |
| **200-$LC_{dc}$** | 3.7 | 3.5 | 0.5 |
| **200-$LCC_{dc}$** | 1.1 | 1.3 | 0.2 |
| **400-$C_{dc}$** | 17.1 | 12 | 2.5 |
| **400-$LC_{dc}$l** | 5.4 | 2.4 | 0.5 |
| **400-$LCC_{dc}$** | 12 | 3.7 | 0.8 |
| **200-Gelatin$_{es}C_{dc}$** *(control experiment)* | 35.8 | 14.9 | |
| **200-Gelatin$_{es}$** *(control experiment)* | 3.7 | 1.7 | |

**Table S1:** Mean Young modulus values for all ECM models measured by peak force tapping AFM-nanoindentation, with standard deviation and standard error of the mean. Force-distance curves were measured in at least 8 positions for 3 different areas of each ECM model.

| | | | **p values** |
|---|---|---|---|
| ***Comparison of protein composition for a given process*** | ***es*** | **$C_{es}$** *vs* **$LC_{es}$** | 0.99993 |
| | | **$C_{es}$** *vs* **$LCC_{es}$** | 0.93296 |
| | | **$LC_{es}$** *vs* **$LCC_{es}$** | 0.99627 |
| | ***dc*** *200 µm* | **200-$C_{dc}$** *vs* **200-$LC_{dc}$** | 1.89374E-7 |
| | | **200-$C_{dc}$** *vs* **200-$LCC_{dc}$** | 0 |
| | | **200-$LC_{dc}$** *vs* **200-$LCC_{dc}$** | 0.58751 |
| | ***dc*** *400 µm* | **400-$C_{dc}$** *vs* **400-$LC_{dc}$** | 9.23236E-8 |
| | | **400-$C_{dc}$** *vs* **400-$LCC_{dc}$** | 0.14555 |
| | | **400-$LCC_{dc}$** *vs* **400-$LC_{dc}$** | 0.01589 |
| ***Comparison of process for a given protein composition*** | *C* | **$C_{es}$** *vs* **200-$C_{dc}$** | 0 |
| | | **$C_{es}$** *vs* **400-$C_{dc}$** | 7.08558E-8 |
| | | **400-$C_{dc}$** *vs* **200-$C_{dc}$** | 4.30343E-9 |
| | *LC* | **$LC_{es}$** *vs* **200-$LC_{dc}$** | 0.85009 |
| | | **400-$LC_{dc}$** *vs* **200-$LC_{dc}$** | 0.98153 |
| | | **$LC_{es}$** *vs* **400-$LC_{dc}$** | 0.99998 |
| | *LCC* | **$LCC_{es}$** *vs* **200-$LCC_{dc}$** | 0.00225 |
| | | **$LCC_{es}$** *vs* **400-$LCC_{dc}$** | 0.36379 |
| | | **400-$LCC_{dc}$** *vs* **200-$LCC_{dc}$** | 2.82647E-8 |

**Table S2:** Mean Young modulus values measured by AFM nanoindentation: analysis of variance (ANOVA) with the Tukey's Multiple Comparison test. P values < 0.05 were deemed significant.
